# A Cryo-EM Grid Preparation Device for Time-Resolved Structural Studies

**DOI:** 10.1101/563254

**Authors:** Dimitrios Kontziampasis, David P. Klebl, Matthew G. Iadanza, Charlotte A. Scarff, Florian Kopf, Frank Sobott, Diana C.F. Monteiro, Martin Trebbin, Stephen P. Muench, Howard D. White

## Abstract

Structural biology generally provides static snapshots of protein conformations that can inform on the functional mechanisms of biological systems. Time-resolved structural biology provides a means to visualise, at near-atomic resolution, the dynamic conformational changes that macromolecules undergo as they function. Recent advances in the resolution obtainable by electron microscopy (EM) and the broad range of samples that can be studied makes it ideally suited to time-resolved studies. Here we describe a cryo-electron microscopy grid preparation device that permits rapid mixing, voltage assisted spraying, and vitrification of samples. We show that the device produces grids of sufficient ice quality to enable data collection from single grids that results in a sub 4 Å reconstruction. Rapid mixing can be achieved by blot and spray or mix and spray approaches with a delay of ~10 ms, providing greater temporal resolution than previously reported approaches.

## Introduction

Our fundamental understanding of biological processes is often underpinned by structural biology, which ultimately may assist our ability to design tailored medicines through structure-based drug design^1^. Advances in technology are allowing us to determine the structure of more complicated and challenging systems. Yet, these methods typically yield structural snapshots of single states of otherwise dynamic protein systems. Better understanding of molecular mechanisms in biology requires time-resolution, which is restricted by current technology. X-ray Free Electron Laser technology can resolve side chain movements during catalysis on the ps to minute timescale, but its use is limited to samples that form ordered micro-crystals^2,3^. Cryo-electron microscopy (cryo-EM) is not constrained by the crystal lattice and allows for structure determination of large non-symmetric macromolecular complex structures in solution, to near-atomic resolution^4^. Cryo-EM has been used to visualise large structural changes in proteins and protein complexes on the millisecond (ms) timescale^5^ and used to determine the structure of multiple conformational states exhibited by numerous macromolecular systems, such as the ribosome and rotary F-ATPase^6,7^. However, it has proven difficult to trap different kinetic sub-states and, although computational sorting can provide different conformations, it does not inform on the order of catalysis, or the lifetime of the intermediates^8^.

The recent renaissance in cryo-EM has been fuelled by rapid developments in microscopes, direct electron detectors, and ever more sophisticated data processing algorithms, but a current bottleneck still resides within the sample preparation procedure, which has changed little over the last 30 years. There have been recent advances in grid production technology such as spotiton™ and vitrojet™, which show great promise in producing high quality and consistent ice but they are not currently capable of time-resolved applications^9^.

Time-resolved cryo-EM (**TrEM**) is an ideal method to study conformational changes occurring on the μs to ms time-scale^10^, providing snapshots of conformational states at sub nm resolution that can be put in order to understand mechanisms on a broad range of specimens. TrEM experiments have previously been demonstrated using a blot and spray approach; applying the protein of interest to an EM grid and pre-blotting before spraying substrate to initiate a reaction, followed by rapid plunging into liquid ethane, to vitrify and stop the reaction at a specific time-delay. This was demonstrated by the ground-breaking work on the acetyl choline esterase receptor^11^. Alternatively, in a ‘mix and spray approach substrate and protein can be pre-mixed in a microfluidic chamber, before directly spraying onto a fast-moving EM grid, which is plunged into liquid ethane^12,13^. This technique has shown great promise for studies of ribosome mechanics.

Here we report a modified device, based on previous designs, which provides a step-change in our capabilities for TrEM. This is the first apparatus that permits both blotting and spraying of EM grids, as well as rapid mixing and that takes advantage of the benefits of voltage-assisted spraying (typically 5 kV). The resulting grids are of sufficient quality for sub 4Å resolution structure determination and allow rapid mixing and freezing in the ms time-frames (>10ms). These capabilities will allow us to determine different conformational states of protein complexes with a time resolution not accessible through conventional EM grid preparation methods.

## Methods

### Sample preparation

Apoferritin from equine spleen was obtained from Sigma Aldrich (A3660), and dialysed into the target buffer (20 mM Hepes, 150 mM NaCl pH 7). *E. coli* ribosomes were prepared in 50 mM Hepes pH 7.5, 100 mM KAc, 8 mM MgAc_2_. Porcine ventricular thin filaments (PV-TF) were prepared as previously described^15^ and diluted to 10 μM in 10 mM Mops pH 7, 50 mM KAc, 3 mM MgCl_2_, 1 mM EGTA prior to use. For the mixing/spraying experiment with apoferritin and thin filaments, 30 μM apoferritin and 10 μM thin filaments were used and mixed in a 1:1 ratio. For the actomyosin dissociation experiment, a solution of 20 μM thin filaments with 24 μM myosin S1 in 10 mM Mops pH 7, 50 mM KAc, 3 mM MgCl_2_, 1 mM EGTA was rapidly mixed (1:1) with buffer containing 0, 0.016, 0.16 or 1.6 mM ATP (making final ATP concentrations of 0, 8, 80, 800 μM). G-actin used for the blot/spray experiment was polymerised according to a protocol by Professor Peter Knight (personal communication).

### Grid preparation

Before spraying/plunge freezing grids, all syringe pumps were initialised and the generated spray was examined visually. Compressed N_2_-gas was bubbled through water to increase humidity in the spraying chamber and avoid evaporation of the spray in flight. The position of the sprayer was adjusted, using a trial grid to ensure it was aligned. A styrofoam system from a Mark IV vitrobot was used to provide a reservoir of liquid nitrogen and to house the liquid ethane. Valves controlling plunger speed and N_2_ flow were adjusted to provide the desired flow rate and plunger speed. The tubing and syringes of the plunge freezing apparatus was washed with 3 x 50 μL ddH_2_O, followed by 3 x 50 μL of the respective buffer and loaded with 33 μL protein solution (the dead volume between the valves and the sprayer is 25 – 35 μL). Cryo-EM grids (Quantifoil R2/1 Cu mesh 200-300 or Quantifoil R1.2/1.3 Cu mesh 300) were glow discharged with air plasma in a Cressington 208 carbon coater with glow discharge unit for 90 s (10 mA/0.1 mbar air pressure) and used within 30 min to avoid hydrophilic recovery of the grids. For grid preparation negative pressure tweezers holding a grid were mounted into the plunging arm of the instrument and the environmental chamber of the instrument was closed allowing the humidity to reach ≥80-90%. 33 μL of protein solution was loaded into a 100 μL Hamilton syringe. The ethane cup, was placed in the target position and the high voltage was turned on. A software-controlled system was used to control the flow rates and timing. Typically, the solution was sprayed at 8.3 μL/s for 3.5 s (to stabilize the spray) before plunging through the spray. A pressure of 0.5 – 2.5 bar was applied to the piston causing the grid to accelerate and reach the spray in less than 0.1 s (distance: 4 cm). The grid passed through the spray, accelerated further over the remaining 3 cm to the ethane surface and the movement terminated by a mechanical stop leaving the grid ~1cm below the surface of the liquid ethane.

The grid was then transferred into liquid N_2_ and stored until screening. For the mix/spray-technique, two syringes were used, one with each reactant and the flow rate was reduced accordingly, to 4.2 μL/s for each syringe resulting in the same overall flowrate. For the blot and spray-technique 3 μL of the first protein sample was placed onto the grid, which was subsequently blotted with two strips of Whatman 43 filter paper for 4 s. After blotting the blotter was retracted and the grid plunged through the spray (which had been initiated during the blotting step to ensure stabilisation) into liquid ethane. Upon completion of all experiments the syringes and tubing were washed with buffer, water and 20 % ethanol aqueous solution.

### Data collection and processing

All cryo-EM imaging was done on a FEI Titan Krios microscope equipped with a Falcon III direct electron detector operating in integrating mode (Astbury BioStructure Laboratory). The main data acquisition and processing parameters are listed in Table S1. All processing was done using RELION2.1^14^ and RELION 3 beta^15^. For all three datasets, the processing was done as follows: Micrographs were motion corrected with RELION 3’s implementation of MotionCor2^16^ and the CTF for each micrograph was estimated using GCTF^17^. For apoferritin and ribosome, references for automated particle picking were generated from class averages from a small number of manually selected particles. The results of the template based automated picking were manually inspected. 2D classification was used to select particles with high resolution information, which were taken forward to generate an initial 3D reconstruction. After two rounds of CTF-refinement and Bayesian particle polishing the final reconstruction was obtained. The presented structures were filtered by local resolution.

For the ribosome dataset, 3D classification with 6 classes was used to sort out 50S subunits reducing the particle number from 47,866 to 34,010. For the thin filament dataset, the RELION tools for helical processing were employed.^18^ All filaments were manually picked. As an initial model, either a 60 Å lowpass filtered structure of actin (PDB: 5mvy), or a featureless cylinder were used. Both gave nearly identical results with the lowpass filtered PDB model leading to slightly higher resolution. CTF refinement was not found to be beneficial in this case. The final reconstruction was subjected to 3D classification with 8 classes to extract a tropomyosin-containing structure (9496 particles) which after 3D refinement had a resolution of 10.4 Å. Helical symmetry for both structures was applied using the refined values for twist and rise with the relion_helix_toolbox program. FSC was determined by the two half map gold standard method for which curves are shown in Supp. Fig. S1.

### Mixer Design

The mixer/sprayer was constructed from standard HPLC fittings and tubing, and the air tip from a 10 μl Gilson pipette tip (Supp. Fig. S2). The spray tip (Supp. Fig. S2 A) was constructed from a 2-3 mm length of 150/40 μm polyimide coated quart tubing (Molex) glued into 360/180 μm tubing with polyacrylate glue. The tubing was sealed into 1/16” O.D. FEP tubing (Upchurch) with a 0.015” I.D. The spray tip was positioned 0-0.5 mm past the end of the air tip. Connection to the high voltage was via a short piece of .007” platinum wire used to make contact with the solution via a “T” connector with 1/16” O.D., through 0.01” I.D. FEP tubing. The voltage of the HV supply could be varied from 2 to 10kV by varying the input from a low voltage DC power supply (Celex BPS1510) to an EMCO Q101N-5 HV converter. The High Voltage (HV) was measured from the voltage across the 100KΩ section of a 100M-100KΩ voltage divider circuit using a DVM (Radio Shack). The valves and pistons of the syringe pumps and forceps holding the EM grid were all grounded to prevent possible damage from stray HV.

For the mixing units (Supp. Fig. S2 B), two concentric tubes (360/200 μm and 100/165 μm) (Molex) were used so that contact of the two solutions would not occur until just prior to spraying. FEP tubing (.007” I.<u>D</u>. 1/16” O.D.) was used to seal the inner 165 μm capillary. The interior tubing was positioned to be 100-200 μm from the beginning of the spray tip to minimize the dead time. The mixer’s geometry approximates that of a double back-to-back “T” mixer.

## Results

The modified setup described in this work (Fig. 1) is based on a previous design^10,13^, to which a number of significant adaptations have been applied. To improve spray distribution and permit a finer aerosol of droplets a 5kV potential is used for voltage-assisted spraying, which has significant advantages over standard spraying using only air pressure. With this approach, the size of droplets produced can be more stringently controlled so as to obtain optimal ice thickness whilst requiring less sample. Droplets produced by voltage assisted spraying are highly-charged, promoting their self-dispersion and preventing coalescence^19^. Additionally, a humidity-controlled chamber is used to provide reproducible conditions and ensure that the micro-droplet spray does not evaporate *en-route* to the EM grid. The system is operated through computer-controlled syringe drivers (Kloehn 50300 series) that control the timing of and flow of the spraying, and additionally provide software control of sample blotting and plunging. The basic design has been described in detail in a previous review^10^.

**Figure 1.**
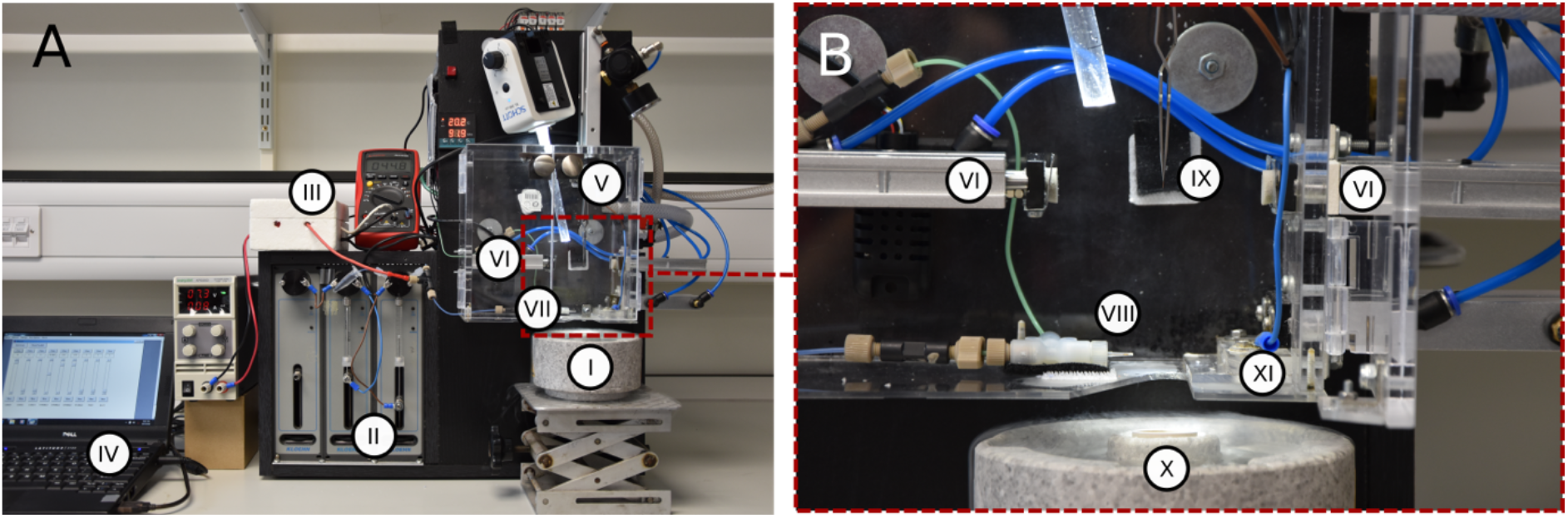
Current setup of the TrEM apparatus. (A) Overview of the complete apparatus showing the styrofoam freezing cup which houses the liquid ethane (I); syringe pumps (II); High Tension (HT) voltage module (III); computer controller (IV); forceps on plunger (V); blotting arms (VI); and sprayer (VII). (B) Zoomed-in view of the spray chamber showing the spray nozzle (VIII); blotting arms (VI); forceps with grid (IX); ethane cup within Styrofoam liquid nitrogen holder (X); and a port that opens just prior to the grid plunge and limits the exposure the liquid ethane to the humid air in the chamber (XI).

As in the case of traditional blotting, plasma treatment of the grid’s surface is an essential step to reduce the inherent hydrophobicity of the carbon coated grids; otherwise very few, if any, droplets adhere to the grid. In addition to the time of mixing, the plunge speed of the grid was used to alter the delay time and time-resolution of the experiments. In our apparatus, we use dual chamber air pistons (Fig. 1) to blot the grids (VI), accelerate the grid into the liquid ethane (I), and open and close the port opening (XI). To better define the parameters and speeds at which grids could be plunged, a range of air pressures (0.5 to 2.5 bar) were used, producing plunge velocities typically at 3 m/s (Supp. Movie 1).

We initially investigated the ability of the TrEM set-up to produce grids with ice of sufficient quality and to collect cryoEM data capable of generating high resolution structures. Three different model systems (apoferritin, ribosome, and thin filaments) were used to investigate the ability to produce samples when spraying on a fast-moving vertical plunger. Data for each system were collected from single grids, using a Falcon 3 detector running in integrating mode on an overnight run (~12 hrs). From overview images, we observed ~ 15-20 % of the grids’ surface area covered with droplets. Although many droplets produced ice too thick for data collection, the grids showed many areas of ice that were suitable for imaging and subsequent high-resolution structure determination (Supp. Fig 3). For apoferritin, a total of 1772 micrographs were collected, with the resulting reconstruction producing a global resolution of 3.6 Å (Figure 2A,B). For the *E. coli* ribosome, 1494 micrographs were collected, with 34010 particles contributing to a final reconstruction that had a global resolution of 4.3 Å (Figure 2C,D) and showed a wide sampling of angular orientations (Figure S4). Porcine cardiac thin filaments were studied to investigate the applicability of the apparatus on long thin filamentous particles. The resulting grids showed good distribution of the specimen within the holes, with no signs of damage in terms of filament length (Figure 2E,F). The resulting reconstruction of the actin core of the thin filaments was resolved to 5.6 Å with the entire thin filament having a lower resolution (10.4 Å, Figure S4), due to heterogeneity of the attached tropomyosin. The results confirmed that this spraying apparatus is capable of producing ice quality sufficient for high resolution cryo-EM data collection and that it can be used as an alternative to blotting for sample preparation.

**Figure 2.**
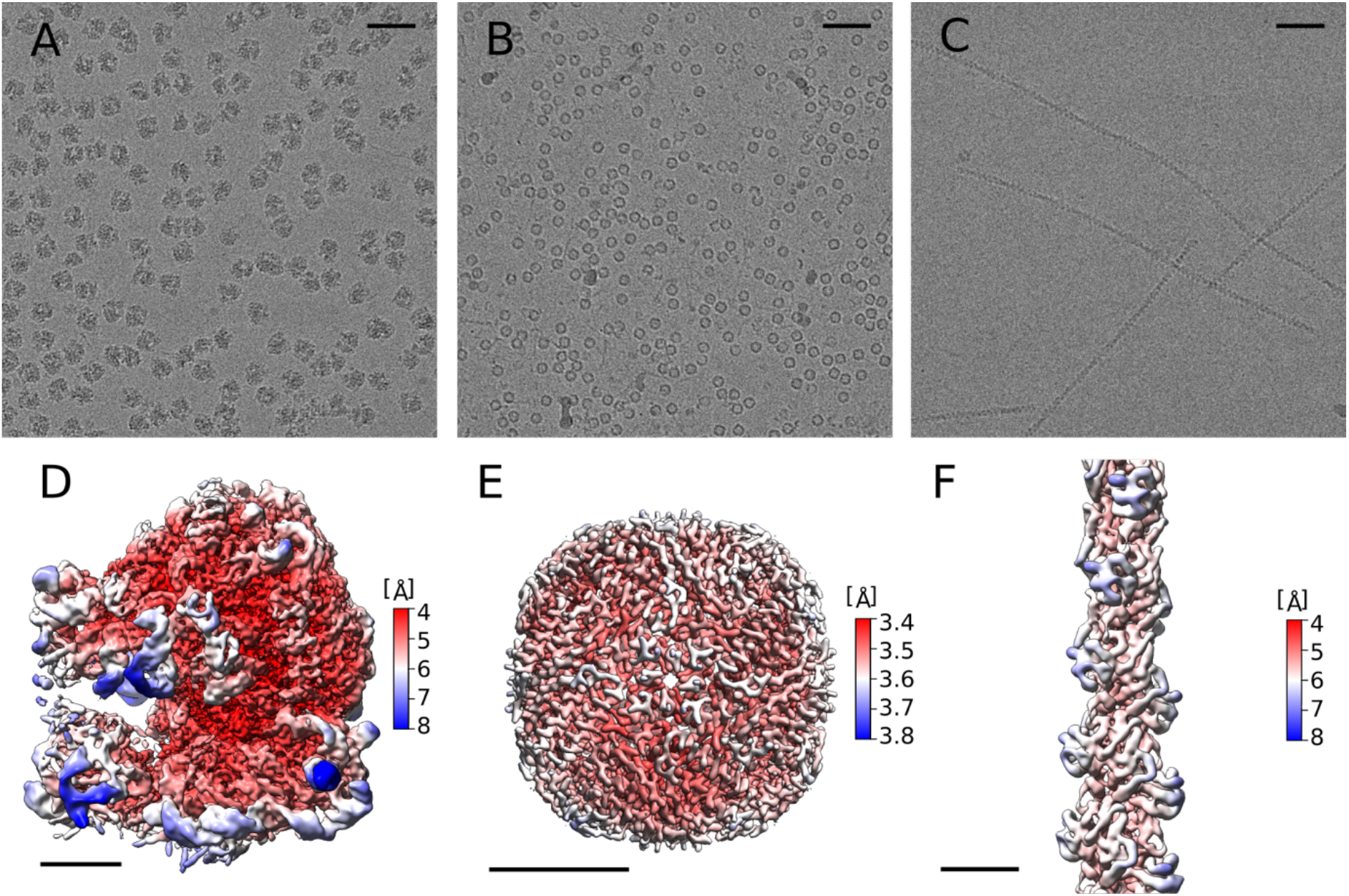
Structures of the 3 model systems from grids prepared on the TrEM setup. To test our ability to make high quality EM grids by spraying proteins on a fast moving plunging grid, three samples were tested and data are shown with representative micrographs and 3D reconstructions: *E. coli* Ribosome (0.72 μM in 50 mM Hepes pH 7.5, 100 mM KAc, 8 mM MgAc_2_) (A, D); apoferritin (30 μM (24mer) in 20 mM Hepes pH 7.5, 150 mM NaCl) (B,E); and porcine thin filaments 5 μM (actin monomer) in 10 mM Mops pH 7, 50 mM KAc, 3 mM MgCl_2_, 1 mM EGTA) (C,F). Scale bar in A, B and C represents 50 nm. Scale bar in D, E and F represents 5 nm.

### Estimation of plunging and droplet speed

In a mix-and-spray type of experiment, the total amount of time the mixed substrates interact can be calculated if we know the speed of the plunger arm, the distance travelled and the speed of the droplets after leaving the mixer/sprayer. The speed of the pneumatic plunging arm was determined using a linear potentiometer. Although it should be noted that the plunger continuously accelerates as the grid moves towards the ethane container. Therefore, the measurement is a single measure of velocity at the position of the spraying nozzle, which slightly underestimates the overall plunging speed. With a constant distance of 3 cm between the central point of the spray cone and the ethane surface, we calculate a time between 30 and 10 ms (for a pressure range of 0.5 – 2.5 bar) for the grid to reach the ethane once the sample has been applied (Fig. 3 A). Droplet speeds were calculated based on high speed video recordings of sprayed droplets at the tip of the nozzle, as well as at a distance of 4 mm from the nozzle (green/blue square in Fig. 3B). Both yield droplet speeds of greater than 4 m/s (Fig. 3C). We also observed acceleration of the droplets as they moved away from the sprayer. With a tip/grid-distance of 6 – 15 mm, this equates to a time-of-flight of ≤ 4 ms for the slowest droplets and the longest distance. Higher plunge speeds result in fewer number of droplets adhering to the grid, with the shortest possible delay on the current setup equating to ~11 ms from the time the droplet leaves the spray nozzle to it being vitrified on the grid (with a typical drop speed 6m/s a 6mm sprayer to grid distance and 3cm spray to ethane distance).

**Figure 3:**
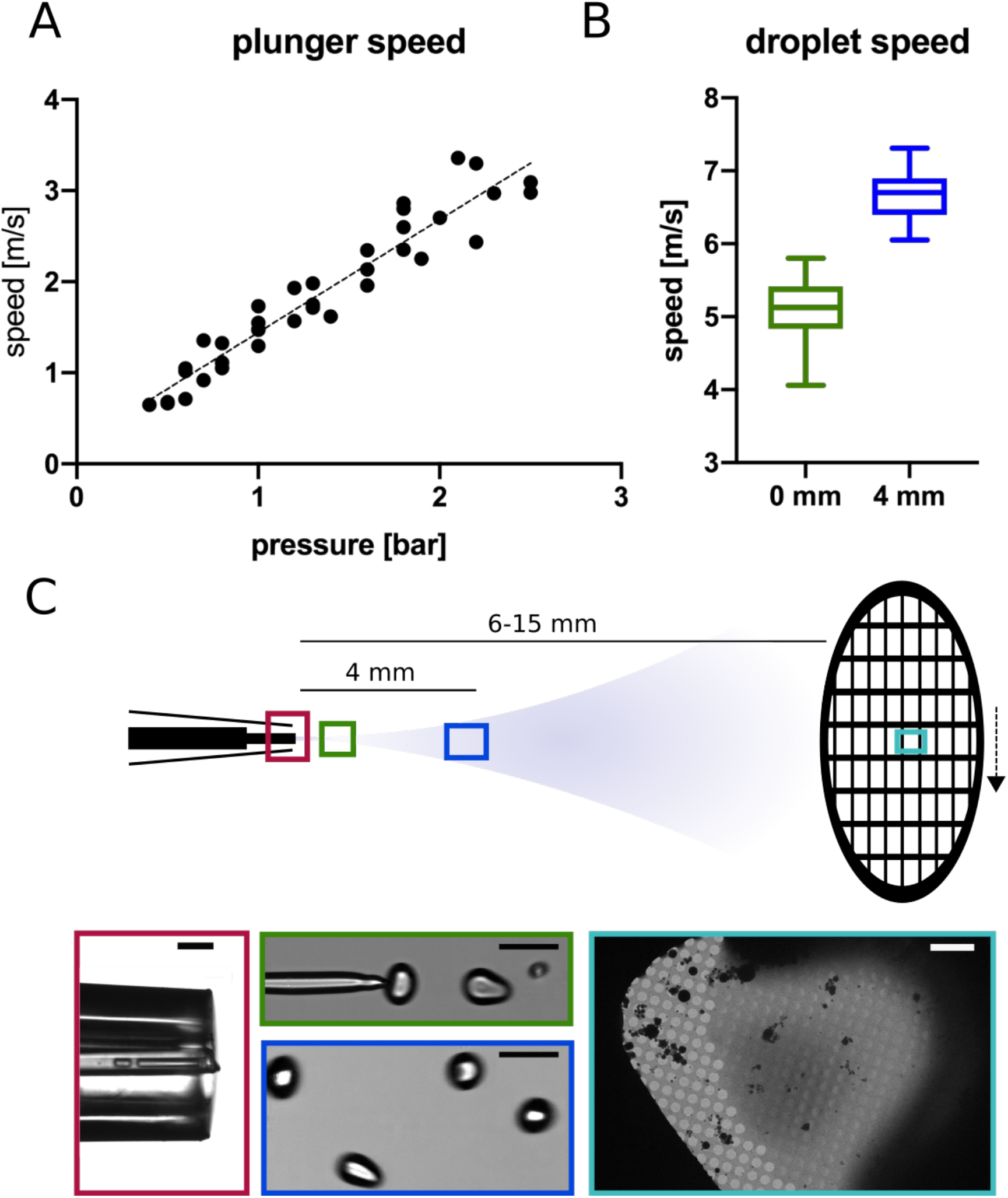
Measuring plunger and droplet speeds. (A) The relationship between pressure and plunger speed, linear fit: speed [m/s] = ~1.5 pressure [bar]. (B) A boxplot of droplet speeds with 10 different droplets tracked over at least three frames for each position. (C) Microscopic images of the spray tip (red, scale bar 200 μm), breakup point of the liquid jet (green, scale bar 100 μm), 4 mm away from the capillary tip (blue, scale bar 100 μm) and grid square of a vitrified grid in the electron microscope (cyan, scale bar 10 μm).

In order to examine the applicability of the new apparatus for the preparation of time-resolved samples, we validated the mixing capability using several different approaches. The first was to conventionally blot the grid containing the first protein and subsequently pass the blotted grid through a voltage assisted spray containing the second protein (Supp. Movie 2). By removing the dead time associated with the mixing chamber and in-flight time of the droplets the time-delay between protein deposition on the grid and freezing is ~ 10 ms, and is dependent only on the distance between the sprayer and liquid ethane (3 cm), and plunge speed (≤ 3 m/s). Actin filaments were pre-blotted on the grid (blot time 4 s) and then passed through a voltage assisted spray of apoferritin. Good quality and clear colocalization could be detected for the samples (Fig. 4A). The second approach was to mix apoferritin and thin filaments within the capillary tube just prior to voltage assisted spraying with the resulting ice showing clear mixing of both samples in all areas studied (Supp. Movie 3 & Fig. 4B). The added delay time, resulting from the additional volume prior to spraying is 1-2 ms, and time of flight from the spray trip to the grid was measured to be less than 4 ms resulting in an overall time delay ~ 15 ms. Although both of these approaches provided promising results and showed clear colocalization, the next step was to provide clear evidence of mixing. To achieve this, we rapidly mixed and vitrified two samples by mixing actomyosin-S1 with varying concentrations of MgATP, which dissociates the myosin-S1 from the actin with a second order rate constant of 10^6^ M^−1^s^−1 20^. The extent of dissociation of S1 from actin after ~15 ms of mixing of 0, 8, 80 and 800 μM MgATP is predicted to be 0, 10, 70 and >99%. By maintaining the plunger speed in a way that the TrEM setup was working with a ~15ms delay, the resulting filament decoration seen in the microscope was consistent with that expected, demonstrating that the two solutions were mixed for ~15 ms, prior to freezing (Fig. 4C). Investigations are currently ongoing to determine the degree of mixing within the chamber, with provisional modelling suggesting most of the mixing may occur within the droplets in-flight. With the increased time of mixing and time-of-flight for the sample, the approximate resulting time-resolution of this approach is on the range of 10-15 ms. Using refined microfluidic mixers, it will be possible to achieve more complete mixing before spraying the mixed sample.

**Figure 4.**
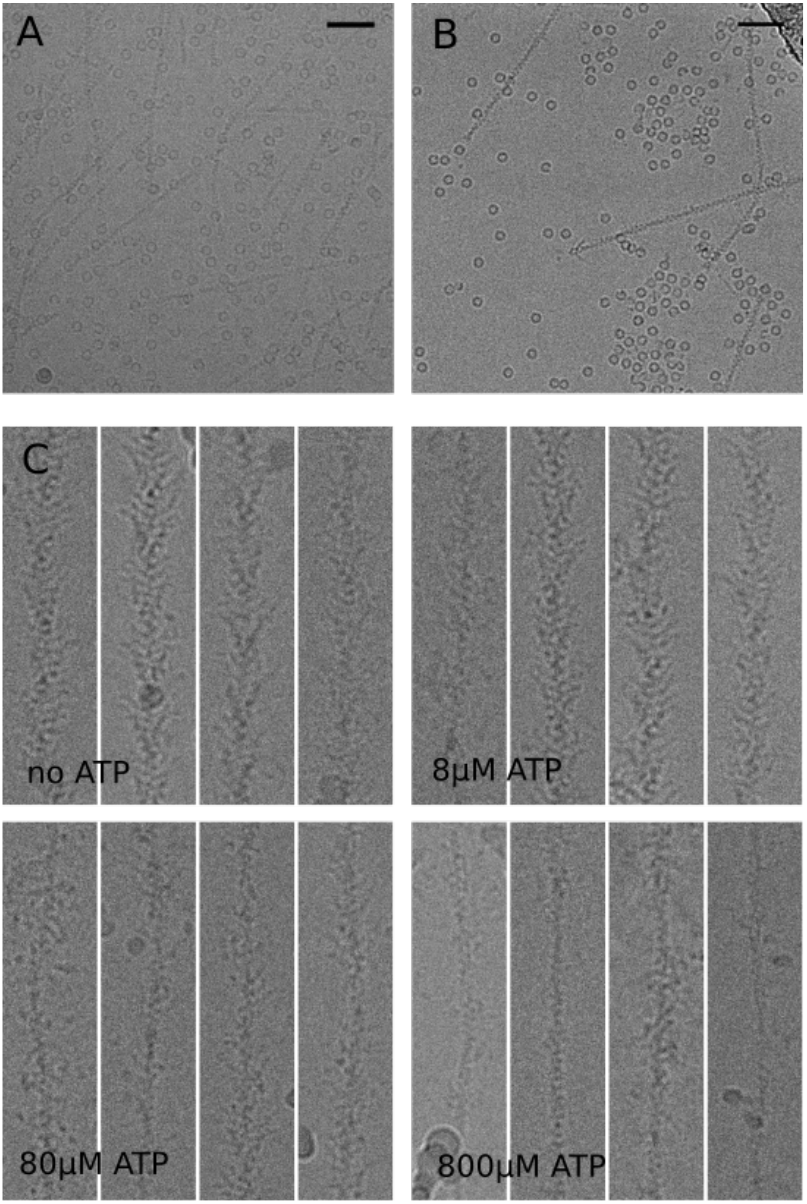
Rapid mixing of samples on the TrEM setup. (A) Representative micrograph with thin filaments blotted and apoferritin sprayed on the subsequent plunged grid. (B) Representative micrograph from the rapid mixing of apoferritin and thin filaments and direct spraying onto the EM grid. (C) Four representative images of Myosin S1 decorated filaments after the rapid mixing (~15ms) of 0, 8, 80 and 800μM MgATP showing decoration consistent with that predicted by kinetic modelling (0, 10%, 70%, 99%).

## Discussion

Electron microscopy has seen significant developments over the last ten years with numerous high-resolution structures of previously intractable protein systems. However, despite the first reported TrEM experiment being conducted in 1991 by the pioneering work of Nigel Unwin, progress has been slow. With the recent developments in cryoEM hardware and software, developing a reliable TrEM setup is becoming a reality. In this work we report an advanced system which can produce cryoEM grids with a minimum time delay between mixing and freezing of 10 ms, faster than the previously reported fastest speed of 24ms^21^. Through three model systems we have shown the capability of producing high quality EM data at a resolution sufficient for resolving side chain density. By using either a blot-and-spray or direct mixing and spraying approach we can produce grids that display clear mixing of the samples, as demonstrated by the dissociation of myosin-S1 from the actin filaments.

Recent studies have suggested rapid vitrification of grids can minimise the interactions with the air-water interface, which is detrimental to the biological sample^22,23^. The methodology reported here is capable of vitrifying grids rapidly and reproducibly, which may alleviate some of the problems associated with interactions at the air-water interface, as well as other problems in conventional blotting systems^24,25^. Ribosome data collected from sprayed grids suggest a more varied distribution of views when compared to sample prepared with the standard blotting method (Fig. S4). This is consistent with the particles making fewer interactions with the air-water interface, however, a full systematic study of this is beyond the scope of this work.

The grid preparation procedure is still a significant bottleneck in cryoEM and has plenty of room for improvements both in terms of reproducibility as well as in the developing of TrEM applications. A number of new approaches have emerged which aim to produce more consistent and reproducible high quality ice, for example spotiton™ and vitrojet™ systems^9^. Here we report a system that addresses a different problem, that of the rapid mixing and trapping of different conformational states to produce cryoEM grids sufficient for high resolution EM structure determination. By using a voltage assisted spray and a rapid mixing unit we can directly spray onto rapidly plunging EM grids or mix and freeze grids quicker than previously reported with a ~10 ms time delay. This integrated apparatus may allow us to open up new opportunities in understanding the mechanism of different protein systems.

## Supporting information

Supplemental Movie 2-Blot and Spray

Supplemental Movie 3-Mix and Spray

## Acknowledgements

This work was supported by a BBSRC grant to SPM and DK (BB/P026397). CAS is funded through a Wellcome Trust ISSF3 Fellowship (204825/Z/16/Z). MGI is supported by MRC grant MR/P018491/1. All EM data were collected at the Astbury Biostructure Laboratory which is supported through the Wellcome Trust (108466/Z/15/Z). EM derived maps have been deposited within the EMDB database (id code EMD-4485 apoferritin, EMD-4488 F-actin, EMD-4487 *E. coli* ribosome) at www.ebi.ac.uk/msd-srv/docs/emdb/. *E. coli* ribosomes were kindly provided by Prof. Neil Ranson, and Mr David Nicholson and G-actin kindly provided by Prof. Michelle Peckham. We would like to dedicate this paper to Prof. John Trinick and his pioneering work on the developments of TrEM applications.

## Contributions

D.K, D.P.K., S.P.M., and H.D.W. developed the apparatus, produced the EM grids and analysed EM data. M.G.I., F.S. and C.A.S. assisted in data analysis and processing. F.K., D.C.F.M, D.P.K. and M.T. conducted high speed imaging of sprayer tips and contributed in the design of the apparatus. S.P.M, H.D.W. conceived and designed the experiments. All authors contributed to writing the manuscript.

---

**Table S1.**
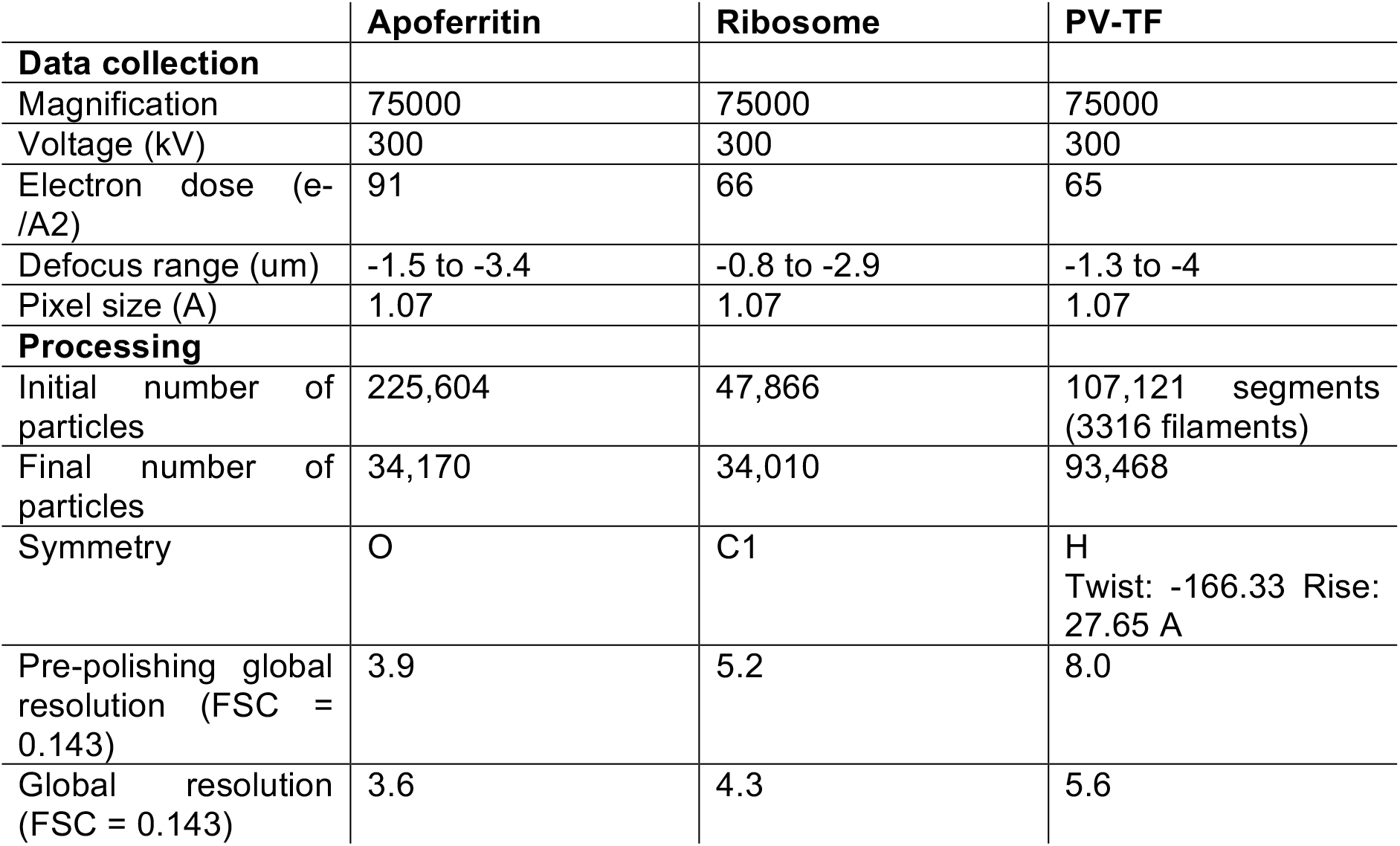
Data collection statistics for the three reported datasets, Apoferritin, ribosome and PV-TF.

**Figure S1:**
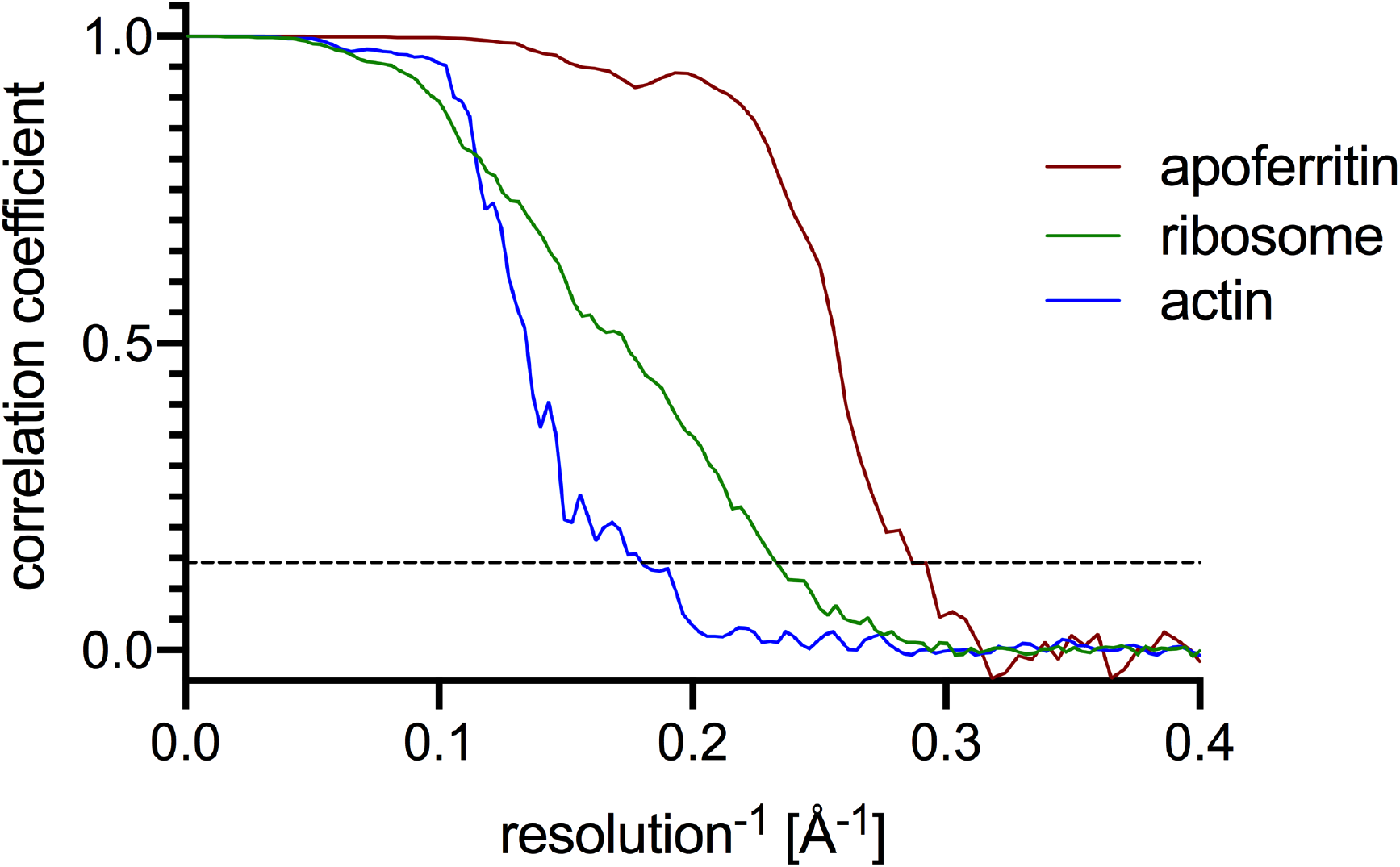
FSC curves. Corrected FSC curves for the final masked maps of apoferritin, actin and ribosome.

**Figure S2:**
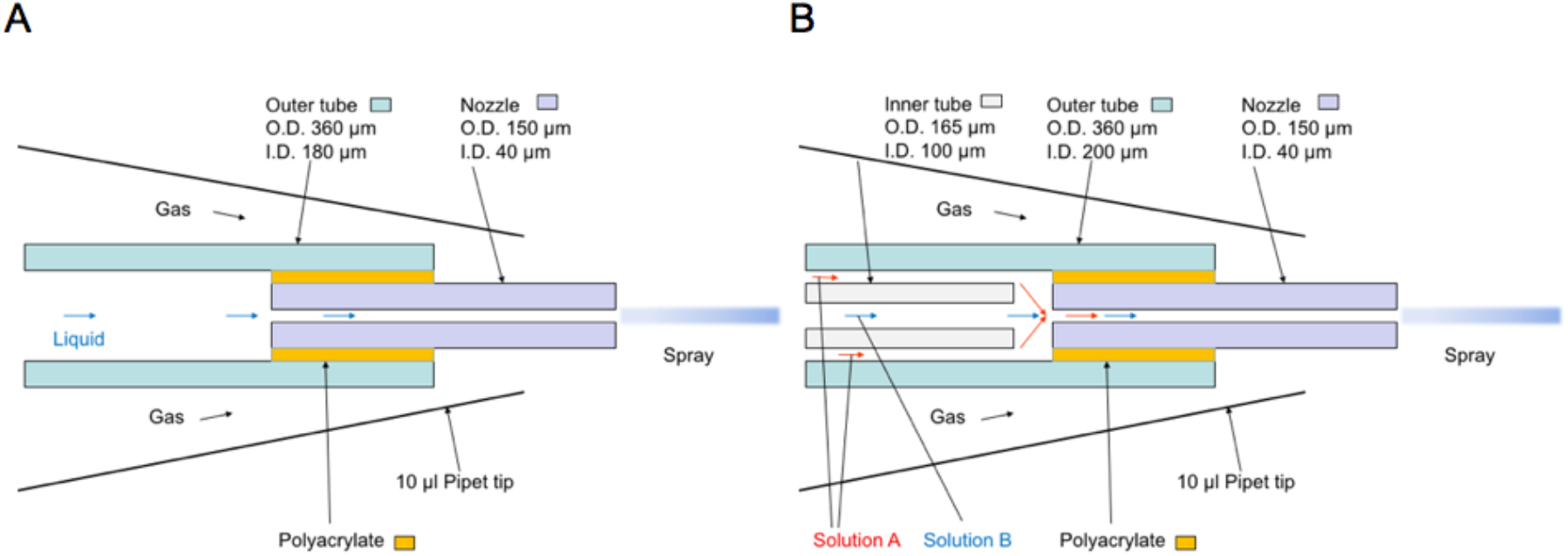
Schematic of the standard spraying unit (A) and the mixing/spraying unit (B).

**Figure S3:**
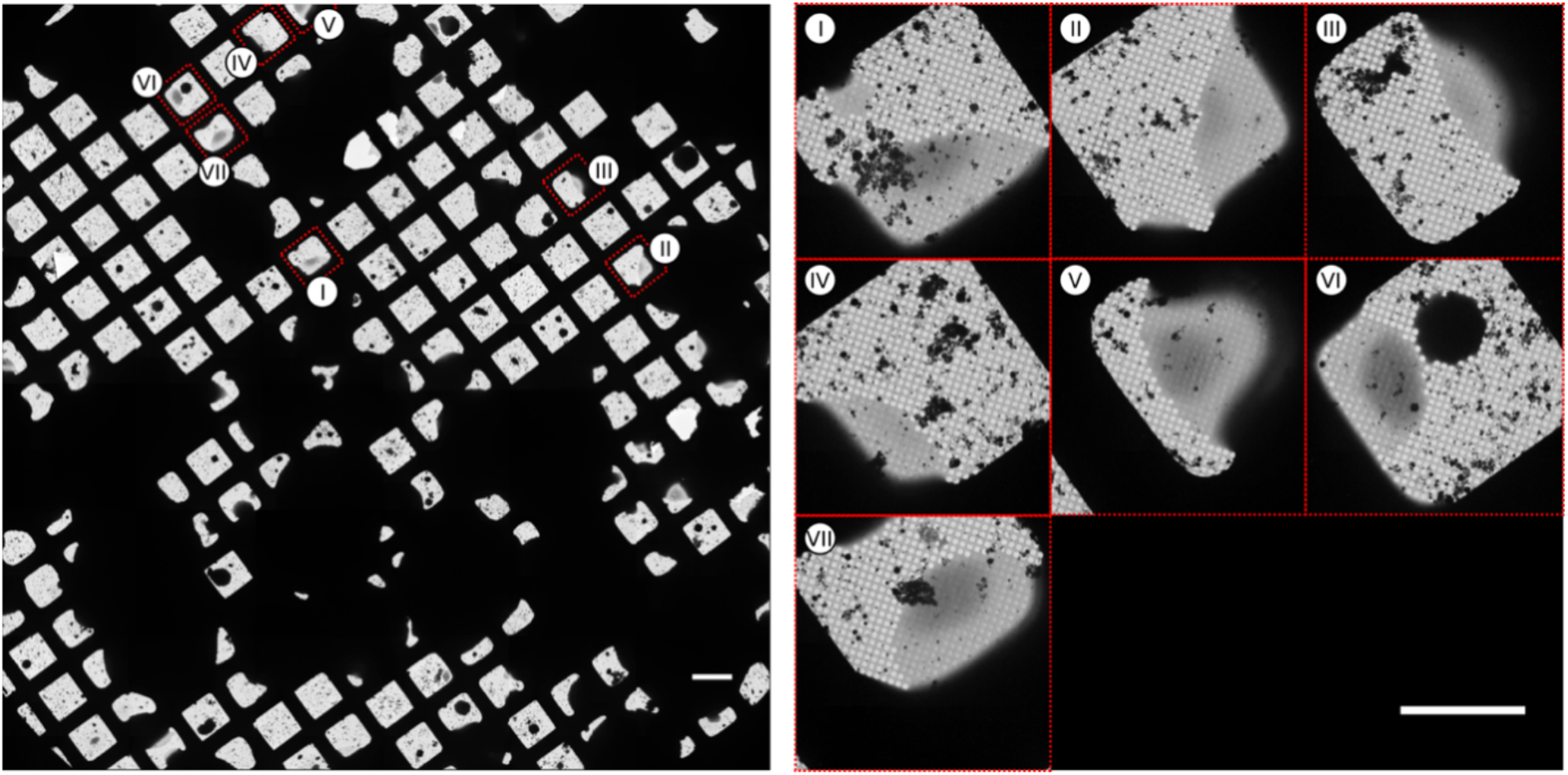
Selection of grid squares for data collection (exemplified for the apoferritin dataset). The image on the left shows a low magnification montage (atlas view) of grid with areas used in data collection highlighted in red (scale bar: 100 μm). The image on the right shows higher magnification micrographs of the selected grid squares with the numbers highlighting the position on the atlas (scale bar: 50 μm).

**Figure S4:**
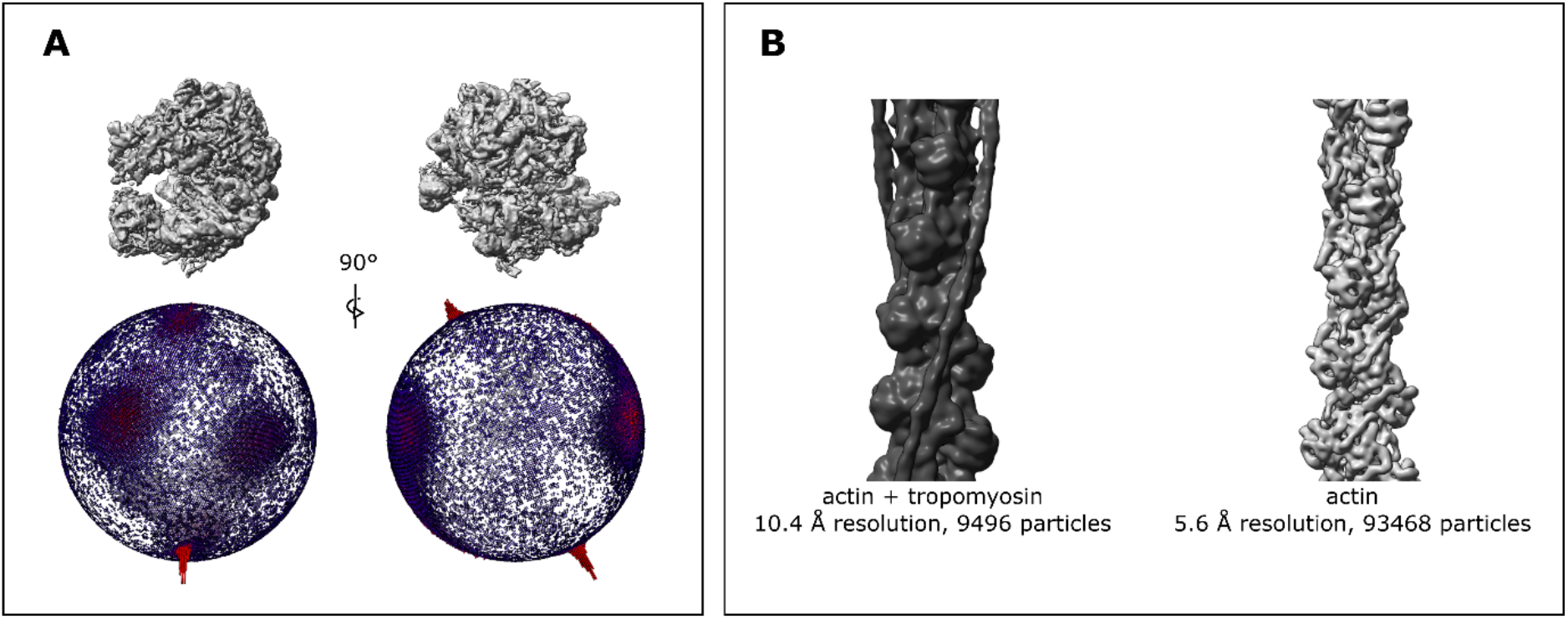
Processing data from sprayed grids. Spread of angular orientation for the ribosome dataset (A) and tropomyosin-containing sub-class in comparison with the consensus structure for the PV-TF dataset (B).

**Movie S1** Real time movie of the TrEM setup in operating in the basic spray mode to generate grids with no pre-mixing or blotting.

**Movie S2** Real time movie of the TrEM setup operating in blot and spray mode. Movie S3 Real time movie of the TrEM setup operating in mix and spray mode.

